# EWS::WT1 Isoform-Dependent Regulation of Neogenes in Desmoplastic Small Round Cell Tumors

**DOI:** 10.1101/2025.09.22.676906

**Authors:** Danh D. Truong, Justin W. Magrath, Kevin Murgas, Jiaqian Fan, Diana Shamsutdinova, Davis Ingram, Alexander Lazar, Sean B. Lee, Joseph Ludwig

## Abstract

Desmoplastic small round cell tumor (DSRCT) is a rare, aggressive sarcoma characterized by the pathognomonic EWS::WT1 fusion protein (FP), an oncogenic chimeric transcription factor (OCTF) resulting from the t(11;22)(p13;q12) translocation. Recent studies have identified “neogenes” (NGs), genes normally silent in normal tissues but transcriptionally activated by OCTFs, as potential tumor-specific markers in fusion-driven cancers. In this study, we investigated the expression and regulation of DSRCT-specific NGs (DSRCT_NGs) using multimodal data across different cohorts of patients, PDX, and cell line data. We evaluated bulk and single-nucleus RNA sequencing of patient specimens from MD Anderson Cancer Center, revealing the robust ability for DSRCT_NGs to distinguish FP-positive DSRCT from samples failing detection of the EWS::WT1 FP. To elucidate the regulatory role of the EWS::WT1 FP in driving NG expression, we performed knockdown experiments in four DSRCT cell lines. This consistently resulted in a reduction of DSRCT_NG expression. Isoform-specific expression of EWS::WT1 in LP9 and MeT-5A mesothelial cells revealed that the E–KTS isoform of EWS::WT1 predominantly drives DSRCT_NG expression. Mechanistically, ATAC-seq and ChIP-seq analyses demonstrated that EWS::WT1 directly binds to accessible chromatin regions near NG transcription start sites, enriched for WT1 motifs and active histone marks. Integration of Hi-ChIP data further revealed that EWS::WT1 facilitates long-range enhancer-promoter looping at DSRCT_NG loci, promoting the expression of nearby genes. Collectively, these findings establish DSRCT_NGs as direct transcriptional outputs of the EWS::WT1 FP and implicate their loci as regulatory regions of the DSRCT transcriptome. Their fusion-dependent expression, chromatin accessibility, and promoter-enhancer connectivity underscore their potential utility as highly specific biomarkers and therapeutic targets in DSRCT.

## Introduction

Several sarcomas have a recurrent gene fusion, which gives rise to an oncogenic chimeric transcription factor (OCTF). These OCTFs act as aberrant transcription factors and have an oncogenic role^1^. Desmoplastic small round cell tumors (DSRCT), first described in 1989 by Gerald and Rosai as a rare but aggressive soft tissue malignancy, are driven by an OCTF^2–4^. DSRCT commonly occurs in adolescents and young adults, whose prognosis remains dismal, with a 5-year overall survival of less than 20%. The OCTF in DSRCT is a result of the t(11;22)(p13:q12) chromosomal translocation between the Ewing sarcoma (ES) breakpoint region 1 gene (*EWSR1*, or *EWS*) and the Wilms tumor transcription factor gene (*WT1*), resulting in the pathognomonic chimeric EWS::WT1 fusion protein (FP) of DSRCT^3,5–8^.

Recently, Vibert et al. showed that the OCTF in fusion-driven sarcomas, like ES and DSRCT, acts as a transcription factor that can drive the expression of sets of “neogenes”^9^. These neogenes (NGs) reside in typically transcriptionally silent regions of the genome and are uniquely activated in the presence of oncogenic FPs such as EWS::FLI1 in Ewing Sarcoma (ES) and EWS::WT1 in DSRCT. Vibert et al. demonstrated that ES-specific NGs (Ew_NGs) are transcribed and translated into neopeptides, suggesting the potential for tumor-specific neoantigens in fusion-driven sarcomas, including DSRCT. Remarkably, NG expression is highly specific to the corresponding OCTF, implying that these genes are direct transcriptional targets of OCTF binding to distinct promoter or enhancer elements. In total, 26 Ew_NGs and 37 DSRCT-specific NGs (DSRCT_NGs) have been identified. However, the regulation and interactions of NGs with EWS::WT1 and DSRCT transcriptome have not been well-characterized. Still, the specificity and restricted expression of NGs underscore their potential utility as biomarkers for diagnosis, measurable residual disease, and molecular subtyping in fusion-driven sarcomas.

In this study, we integrated bulk and single-nucleus RNA sequencing from patient samples, PDX models, and cell lines to demonstrate the strong discriminatory power of DSRCT_NGs in identifying DSRCT. We conducted targeted knockdown experiments to investigate the functional relationship between DSRCT_NGs and the EWS::WT1 fusion protein (FP). Chromatin accessibility profiling revealed specific EWS::WT1 binding sites within NG loci, shedding light on the transcriptional mechanisms driving NG activation in DSRCT. The DSRCT_NG loci function as enhancer-promoter hubs, facilitating EWS::WT1-mediated regulation of proximal gene targets^10^. Collectively, our findings establish DSRCT_NGs as direct transcriptional outputs of the EWS::WT1 FP and highlight their role as key epigenetic regulators of the DSRCT transcriptome.

## Materials and Methods

### Bulk RNA-seq of DSRCT patient specimens

Clinical data, including pathology reports and EMRs, were reviewed for all confirmed DSRCT cases. Tumor specimens were archived in the MDACC Biospecimen Bank or Dr. Ludwig’s lab. Diagnosis was confirmed by expert sarcoma pathologists using clinical features, immunohistochemistry, and cytogenetics to detect the EWS::WT1 fusion (**Supplementary Table 1**). Mapsplice^11^, FusionMap^11^, and TophatFusion^12^ performed fusion call. A specimen was considered fusion-positive when at least two fusion detection tools detected an EWS::WT1 fusion. RNA-seq processing and tissue collection followed previously described methods^13^. Gene expression heatmaps were generated using ComplexHeatmap in R^14^.

Bulk RNA-seq data from DSRCT cell lines (JN-DSRCT-1, BER-DSRCT, BOD-DSRCT, SK-DSRCT2 ± fusion knockdown) and LP9 cells with EWS::WT1 isoform overexpression were obtained from GEO (GSE252051)^15^. Data from MeT-5A cells with EWS::WT1 overexpression came from GSE212976. Gene counts were aligned to a custom transcriptome (GRCh38 + OCTF-driven NG per Vibert et al.) using STAR/RSEM^16^. Differential expression was analyzed with DESeq2^17^.

### Bulk RNA-seq of patient-derived xenografts

The ES, CDS, and DSRCT PDX models were developed in the Ludwig lab (**Supplementary Table 2**). DSRCT-4 is derived from patient T19 and DSRCT-2 is derived from patient T15. Two of the OS PDX (OS1 and OS31) were obtained from the Houghton lab^18^, and one OS PDX (SA98) was developed at the Pediatric Solid Tumors Comprehensive Data Resource Core in MD Anderson Cancer Center (**Supplementary Table 2**).

Total RNA was extracted from frozen desmoplastic small round cell tumor (DSRCT) patient-derived xenograft (PDX) samples using the RNeasy Mini Kit (Qiagen), following the manufacturer’s protocol to ensure high-quality RNA is suitable for sequencing. RNA integrity was assessed using standard quality control measures prior to library preparation. RNA-seq libraries were constructed using the KAPA Stranded RNA-Seq Library Preparation Kit (KAPA Biosystems), which preserves strand orientation and enhances transcriptome coverage. Libraries from individual samples were quantified, pooled in equimolar concentrations, and sequenced on the Illumina HiSeq 4000 platform, generating high-throughput paired-end reads. Sequencing reads were aligned to the human reference genome (hg19) using the GSNAP aligner, which supports spliced alignment and is optimized for detecting fusion transcripts and complex RNA isoforms^19^. Downstream analyses were performed on the aligned data to quantify gene expression and assess NG activation across the PDX cohort. For the calculation of gene expression, raw counts were obtained using featureCounts^20^ and were then normalized by regularized log transformation in DESeq2^17^.

To evaluate specificity of NG expression for DSRCT, CIC-DUX4, and ES (DSRCT-NG, CIC-NG, Ew-NG), Module scores were computed using the AddModuleScore function in Seurat, which calculates the average expression of a gene set relative to control genes. Each signature was scored across all cells and resulting scores were stored in the metadata.

To assess the predictive power of each NG, we randomly split the dataset into training and test groups. For each NG set, we generated matched random gene sets with same number of genes to serve as negative controls. Logistic regression models were trained on the training set using signature scores to predict subtype (e.g., DSRCT vs. others), and performance was evaluated on the test set. Receiver Operating Characteristic (ROC) curves and Area Under the Curve (AUC) values were computed using the pROC package to compare true signature performance against random controls.

### RNAscope in situ hybridization

Formalin-fixed paraffin-embedded (FFPE) tumor blocks from two patients diagnosed with DSRCT were obtained at The University of Texas MD Anderson Cancer Center. For each patient, three consecutive sections were prepared: one with negative control probes, one with positive control probes, and one for the COMET multiomics assay (RNAscope + immunofluorescence + H&E). RNAscope probes targeting four NGs (DSRCT_NG21, NG33, NG6, NG30) were designed by Advanced Cell Diagnostics (Bio-Techne); new probe request number is available upon request. Positive (housekeeping genes) and negative control probes were run separately. Assays were performed on the Lunaphore COMET™ Multiomics platform, at the MD Anderson Flow Cytometry and Cellular Imaging Core Facility, which integrates RNAscope with sequential immunofluorescence, followed by imaging.

Six images were exported as high-resolution TIFF files, imported into Horizon Viewer for background subtraction, and subsequently analyzed in Visiopharm® (Visiopharm, Hørsholm, Denmark). From whole-slide images of 12 × 12 mm², three regions of interest (ROIs) were selected per sample (six ROIs in total). Within each ROI, tumor and stroma regions were delineated by antibody-based thresholding (CACNA2D2 (Santa Cruz, cat no. sc-365911) and pan-cytokeratin (Proteintech, cat no. 26411-1-AP) for tumor; collagen (Cell Signaling Technology, cat no. 66948S), for stroma). RNA puncta were quantified within each region and normalized to area (puncta/µm²). Statistical analysis was performed in Python (v3.9.6) using the Mann-Whitney U test to compare NG probes with positive control probes within each region.

### Cell Culture

DSRCT cell lines, JN-DSRCT-1, BER-DSRCT, BOD-DSRCT, and SK-DSRCT2, were used in this study. All four lines have been previously validated to harbor the characteristic EWS::WT1 fusion and are described in prior publications^15,21–23^. JN-DSRCT-1 was generously provided by Dr. Jun Nishio, while BER-DSRCT, BOD-DSRCT, and SK-DSRCT2 were obtained from Dr. Marc Ladanyi. All cell lines were routinely tested for Mycoplasma contamination via PCR and confirmed negative. To maintain genomic and phenotypic integrity, cells were used for fewer than 10 passages post-thaw. DSRCT cell lines were cultured in DMEM/F12 medium supplemented with 10% fetal bovine serum (FBS; Gibco), 2 mmol/L L-glutamine, 100 U/mL penicillin, and 100 μg/mL streptomycin (Thermo Fisher Scientific). The LP9 cell line, a non-transformed, diploid human mesothelial cell line, was obtained from the NIGMS Human Genetic Cell Repository at the Coriell Institute (catalog number AG07086, Camden, NJ). LP9 cells were maintained in DMEM/F12 medium supplemented with 15% FBS, 2 mmol/L L-glutamine, 100 U/mL penicillin, 100 μg/mL streptomycin, 10 ng/mL epidermal growth factor (EGF; Thermo Fisher Scientific), and 0.4 μg/mL hydrocortisone (Sigma-Aldrich).

### Generation of dox-inducible shRNA and EWS::WT1 overexpression cell lines

To generate stable, doxycycline-inducible knockdown cell lines, we utilized the LT3-GEPIR lentiviral vector, as previously described^24^. Short hairpin RNA (shRNA) sequences against WT1 3′UTR (*5*′ *GCA GCT AAC AAT GTC TGG TTA 3*′) were annealed and cloned into the XhoI and EcoRI restriction sites of the LT3-GEPIR vector. For overexpression studies, EWS::WT1 fusion constructs were cloned into a pCDH lentiviral backbone, as previously established. Lentiviral particles were produced by co-transfecting HEK293T cells with the LT3-GEPIR-shRNA or pCDH-EWS::WT1 constructs along with the ViraPower lentiviral packaging mix (Invitrogen), using Lipofectamine 3000 (Thermo Fisher Scientific) according to the manufacturer’s protocol. Viral supernatants were collected at 48-, 72-, and 96-hours post-transfection and concentrated using Lenti-X Concentrator (Takara Bio). DSRCT and LP9 cells were transduced with lentivirus in the presence of 10 μg/mL polybrene for 16 hours at a multiplicity of infection (MOI) <1. Forty-eight hours post-transduction, cells were selected with puromycin (DSRCT: 0.5 μg/mL; LP9: 1 μg/mL) to establish stable lines. Knockdown efficiency and inducibility were validated by RT-qPCR and Western blotting, both in the presence and absence of doxycycline. For induction of shRNA expression, cells were treated with 1 μg/mL doxycycline.

### ChIP-seq Preprocessing and Visualization

Raw ChIP-seq data in FASTQ format were processed using a custom Bash pipeline. Quality control was performed using FastQC v0.11.9 to assess read quality metrics. Reads were aligned to the GRCh38/hg38reference genome using Bowtie2 v2.4.5 with 28 computational threads. Alignment logs were recorded for each sample. SAM files generated from Bowtie2 were converted to BAM format using Samtools v1.16.1 and subsequently sorted by genomic coordinates using Sambamba v0.8.2. Filtered BAM files were indexed using Samtools for downstream analysis. ChIP-seq data were processed using MACS2 (v2.2.7.1) to identify regions of enriched chromatin marks and transcription factor binding. To visualize ChIP-seq signal intensity across genomic regions, bigWig files were generated and heatmaps were constructed using deepTools (v3.5.4). These visualizations enabled comparative analysis of chromatin features and transcription factor occupancy at NG loci and other regions of interest.

### HiChIP Data Processing and Chromatin Loop Analysis

HiChIP data from the JN-DSRCT-1 cell line were processed using HiC-Pro (v3.1.0) to generate normalized contact maps and identify chromatin loops. Raw paired-end reads were aligned to the human reference genome (hg19) and filtered for valid interaction pairs. Valid interaction pairs were extracted. These interactions were used to identify long-range enhancer-promoter loops, particularly those involving DSRCT_NG loci and EWS::WT1 binding sites.

Loop distances and interaction frequencies were quantified and compared across different categories of loops (e.g., DSRCT_NG–EWS::WT1, intra-DSRCT_NG, and non-EWS::WT1 loops), providing insight into the spatial organization of the chromatin landscape in DSRCT.

To visualize chromatin interactions and regulatory architecture in the JN-DSRCT-1 cell line, we used HiCExperiment^25^ and plotgardener (v1.0.0)^26^ in R. Hi-C contact matrices were imported from HiC-Pro output as a HiCExperiment, specifying the. matrix and .bed files at 20 kb resolution. Looping interactions were annotated using BEDPE files filtered for high-confidence interactions (score > 4).

### Single-nucleus Capture

This protocol was performed as described by Truong et al^27^. Nuclei were isolated from fresh-frozen tissue using the Nuclei EZ Prep Kit (Sigma-Aldrich). Single-nucleus suspensions were stained with 7-AAD in NWRB for 5 minutes on ice for sorting. A BD cell sorter sorted up to 100,000 7-AAD–positive nuclei. Post-sort nuclei concentration and quality were verified microscopically before loading onto a 10x Genomics chip.

### Single-cell sequencing data analysis

Demultiplexed FASTQ files were generated using Cell Ranger mkfastq (10x Genomics). Sequencing reads were aligned to the GRCh38 human reference genome using Cell Ranger v5.0, with the --include-introns option enabled to capture both exonic and intronic reads, allowing for more comprehensive quantification of nascent transcripts^28^. Unique molecular identifiers (UMIs) were counted using the cellranger count pipeline to generate gene-barcode matrices.

Quality control (QC) and normalization followed best practices outlined by the OSCA (Orchestrating Single-Cell Analysis) framework and other established guidelines^29^. QC metrics included total UMI counts, number of detected genes per cell, and the percentage of mitochondrial gene expression. Outlier cells were identified and excluded based on these metrics to ensure high-quality downstream analysis^29^.

Filtered gene-barcode matrices were imported into Seurat v5 for preprocessing and initial analysis. Low-quality cells were removed, and data normalization was performed using SCTransform, a method implemented in the Seurat R package. Clustering was performed using the default Louvain method^30–32^. We generated a two-dimensional embedding using Uniform Manifold Approximation and Projection (UMAP). Heatmaps were created using the ComplexHeatmap and ggplot2 R packages, enabling detailed visualization of gene expression patterns across clusters^33,34^.

ATAC-seq data were processed and analyzed using the Signac package, an extension of the Seurat framework for single-cell chromatin accessibility analysis^30,31^. Initial QC filtering will be applied to retain high-quality single cells filtered for quality by using greater than 1,000 fragments; percent reads in peaks of at least 15%, nucleosome signal less than 10, and TSS enrichment score higher than 2.

Following ATAC QC, data were normalized using the term frequency–inverse document frequency (TF-IDF) transformation, which adjusts for sequencing depth and feature frequency. Dimensionality reduction will be performed using latent semantic indexing (LSI).^35^ We applied the chromVAR inference method for downstream analysis to identify enriched transcription factor binding motifs within accessible chromatin regions. Additionally,

We explored these peak-gene associations by analyzing the co-variation in chromatin accessibility and gene expression across cells through the SHARE-seq framework.

### Data Availability

BAM files for DSRCT patients were previously generated and are available from EGA (EGAS00001004575). ATAC-seq data for DSRCT cell lines (JN, BER) came from NCBI GEO accession GSE248758; A673 from GSE181554; RH4 from GSE116344. RNA-seq, ChIP-seq, and HiChIP data for JN-DSRCT-1 and MeT-5A were sourced from GSE212979. A second ChIP-seq for WT1 binding in JN-DSRCT-1 was acquired from GSE156277. Data for snRNA-seq for PDX were sourced from GSE200529. Data for DSRCT snRNA-seq and single-cell multiome will be available upon accepted publication.

## Results

### Fusion-protein-specific neogene expression in DSRCT

We analyzed a cohort of 40 fresh-frozen DSRCT specimens, each from a patient with a prior diagnosis of DSRCT based on pathologist review of EWS::WT1 FISH staining. Of these specimens, only 23 specimens were considered EWS::WT1 fusion-positive (confirmed by at least two RNAseq-based fusion-detection methods). In contrast, 17 specimens failed fusion calling and were considered EWS::WT1 fusion-negative specimens (**Supplementary Table 1**). These EWS::WT1 fusion-negative specimens likely represent stroma-enriched tissue with few tumor cells. One EWS::WT1 fusion-negative specimen was later reclassified as an ovarian germ cell tumor. In addition, several patients had multiple specimens (T2a-b, T5a-b, T12a-d, T13a-c, T14a-e) resected along with a few matched normal tissue specimens (N6, N8). In some patients with multiple specimens, not all specimens were positive for EWS::WT1, indicating potential stroma-enriched tissue (T12, T23). To clarify the discrepancy, we applied a principal component analysis (PCA) of gene expression profiles and observed the specimens to cluster based on the presence of the FP (**Supplementary** Fig. 1A). Interestingly, we found that T22 did not cluster with fusion-positive specimens even though the EWS::WT1 fusion was called twice in T22. It was called only once in T15, and this specimen clustered with other fusion-positive specimens. Importantly, the rest of the fusion-positive specimens clustered as expected. Differentially expressed gene (DEG) analysis between the fusion-positive and fusion-negative specimens also confirmed the expression of EWS::WT1-specific transcriptomic profile, including previously identified genes like *ST6GALNAC5, CACNA2D2, GAL,* and *GALP* (**Supplementary** Fig. 1B)^36,37^. Although T22 was called a fusion-positive specimen, it did not express DSRCT-specific genes. Gene set enrichment analysis (GSEA) identified upregulated pathways in fusion-positive specimens, including neuronal systems, gap junction assembly, transmission across chemical synapses, and protein-protein interactions at synapses (**Supplementary** Fig. 1C). On the other hand, we found that pathways related to the immune system, inflammation, and adipogenesis were downregulated in the fusion-positive specimens (**Supplementary** Fig. 1D), which echoed our prior findings^37^.

Vibert *et al.* recently identified 37 NGs in DSRCT that are driven explicitly by the EWS::WT1 fusion oncoprotein. We assessed the expression of these 37 DSRCT_NGs in our independent cohort, using publicly available genomic coordinates of the NGs to annotate the NGs in the DSRCT RNA-seq data. We observed that most DSRCT_NGs were consistently expressed across the fusion-positive specimens (**Fig. 1A**). Notably, fusion-negative and matched normal specimens did not express DSRCT_NGs. For instance, in patient T12, who was diagnosed with DSRCT, three specimens were collected, but only one specimen (T12a), called fusion-positive, robustly expressed DSRCT_NGs. In addition, T16 was misdiagnosed with DSRCT and later reclassified as an ovarian germ cell, and this specimen did not express DSRCT_NGs. The two normal tissue specimens, N6 and N8, did not express NGs. Although T22 was called a fusion-positive specimen, it did not express the DSRCT_NGs, consistent with its lack of expression of other DSRCT-specific genes. Likewise, T15 only passed one of the fusion caller methods, but had detectable DSRCT_NGs and clustered with other fusion-positive specimens. These findings indicate that DSRCT_NGs are expressed with high specificity in fusion-positive specimens.

**Figure 1:**
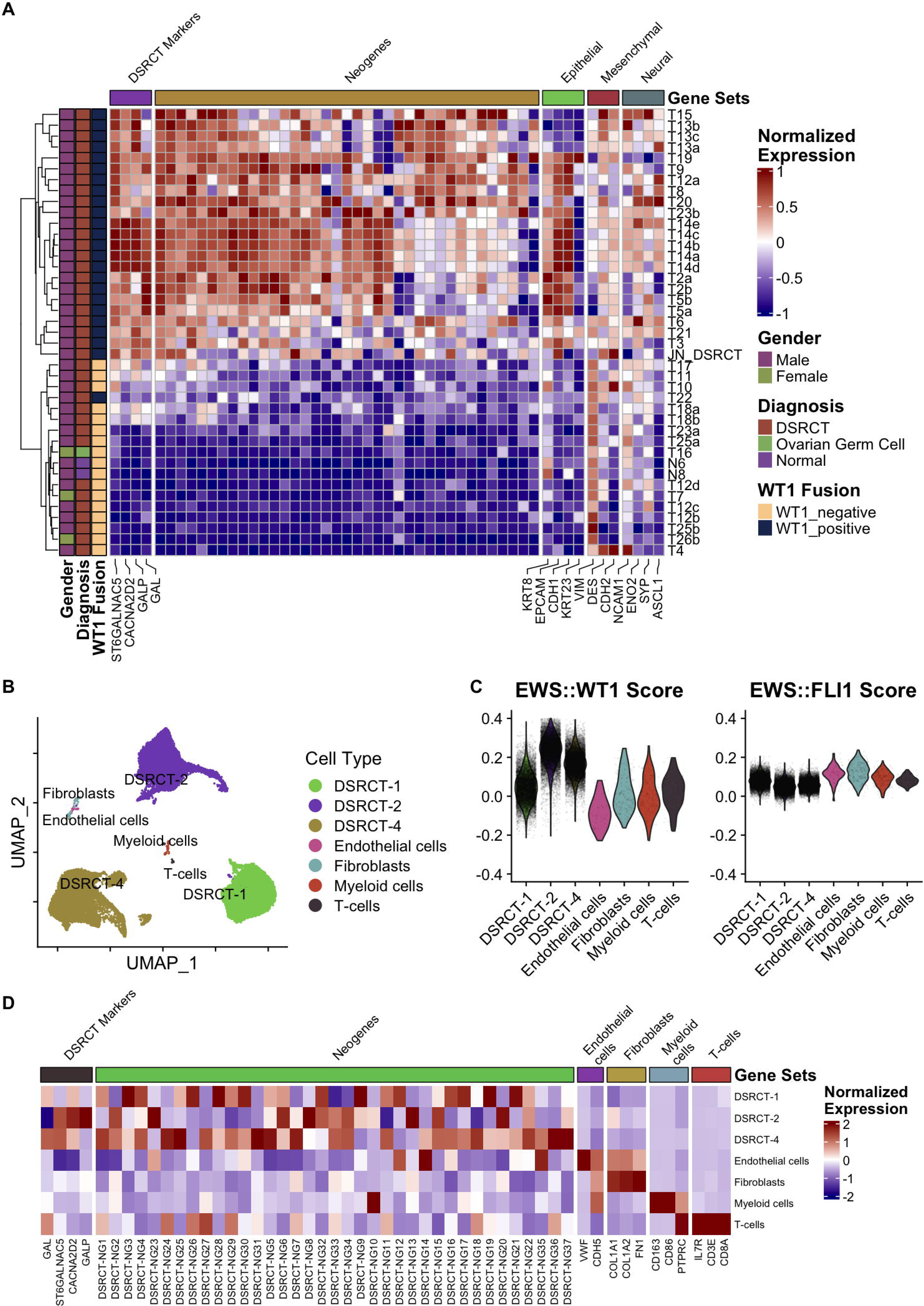
Neogenes are selectively expressed in DSRCT. **A)** Heatmap showing sample-level normalized bulk gene expression of common DSRCT marker genes, epithelial genes, mesenchymal genes, neural genes, and DSRCT NGs. On the left, samples are characterized by gender, diagnosis, and EWS::WT1 fusion. **B)** UMAP projection of single-nucleus data from three DSRCT specimens. Clusters were manually annotated to distinguish tumor and stromal cell classes. **C)** Violin plots of enrichment scores of individual cells for EWS::WT1 and EWS::FLI1 signatures. **D)** Heatmap of average normalized gene expression within identified cell clusters, with associated gene categories annotated.

We also compared the expression of DSRCT_NGs along with DSRCT-specific marker genes and genes associated with multi-lineage expression of epithelial, mesenchymal, and neural markers in DSRCT^38–40^. We observed heterogeneous expression of mesenchymal and neural markers across all specimens, both fusion-positive and fusion-negative. However, DSRCT markers and epithelial-related gene expression, primarily expressed in fusion-positive specimens, correlated with the expression of DSRCT_NGs. Next, we compared the expression of the target genes that were upregulated and downregulated by the EWS::WT1 FP as determined by Gedminas et al^8^. We performed a gene-gene correlation analysis to explore if DSRCT_NGs were correlated with the genes regulated by the EWS::WT1 (**Supplementary** Fig. 2). As expected, we observed a strong positive correlation between the expression of EWS::WT1 upregulated genes and the expression of DSRCT_NGs. Furthermore, the expression of EWS::WT1 downregulated genes was negatively correlated with the DSRCT_NGs. When compared to the expression of DSRCT_NGs, the expression of NGs controlled by other OCTFs was not as strongly expressed or as consistent in DSRCT (**Supplementary** Fig. 3A).

### Single-nucleus expression reveals neogene heterogeneity among specimens

To confirm that DSRCT_NGs were only consistently expressed in the DSRCT cells, rather than stromal components, we explored single-nucleus RNA-seq of three patient samples. We sequenced 40,182 nuclei from these patients. We observed several cell clusters visible on the UMAP, with the largest three consisting of different patient tumor cells (**Fig. 1B**). We annotated the clusters using canonical markers and DSRCT-enriched genes and used the EWS::WT1 signature derived from Gedminas et al. to delineate DSRCT cells (**Fig. 1C**)^8^. An EWS::FLI1 signature comprised of genes that are activated targets of the Ewing-specific EWS::FLI1 signature^41^ was used as a control, which was not enriched in any cluster. We annotated the clusters using canonical markers and DSRCT-enriched genes to identify five subsets of cells: DSRCT cells (within three clusters separated by specimen), fibroblasts, endothelial cells, T-cells, and myeloid cells (**Fig. 1D**). We confirmed that the DSRCT cells highly express *GAL, GALP, CACNA2D2,* and *ST6GALNAC5.* Endothelial cells specifically expressed *VWF* and *PECAM1*. Fibroblasts expressed collagen genes *COL1A1* and *COL1A2.* Immune cells appeared to be a mixture of myeloid cells expressing *PTPRC* (CD45), *CD68*, and *CD163*, and lymphoid cells expressing *CD3E, CD8A*, and *IL7R*. Notably, we observed robust expression of the DSRCT_NGs in only the DSRCT cells and not the non-malignant cells (**Fig. 1D**). To assess the specificity potential of NGs, we applied logistic regression models to classify DSRCT based on NG expression. The dataset was randomly split into training and test sets. Using DSRCT_NGs, the model accurately identified DSRCT cells (AUC = 0.9), outperforming models based on randomly selected genes (AUC = 0.6) or Ew_NGs (AUC = 0.61, **Supplementary** Fig. 3B). This supports the notion that expression of NGs could delineate malignant cells from non-malignant cells within cancers driven by an OCTF^1^.

### Neogene expression is specific for sarcoma subtype in patient and PDX data

Next, to measure NG specificity in a patient-derived setting, we characterized the expression of OCTF-associated NGs in patient-derived xenografts. We investigated snRNA-seq data from DSRCT, Ewing sarcoma (ES), CIC::DUX4 sarcoma (CDS), and osteosarcoma (OS) PDX models. Comparison of pseudo-bulk expression confirmed the expression of subtype-specific marker genes related to the three OCTFs (EWS::WT1, EWS::FLI1, and CIC::DUX4) in their respective PDXs. OS PDXs did not express these gene targets but did express osteoblastic or chondroblastic genes (**Fig. 2A**). We then compared various OCTF signatures based on target genes of each OCTF^8,42,43^. As expected, we observed robust expression of the EWS::WT1 signature only in DSRCT PDXs, the EWS::FLI1 signature in ES PDXs, and the CIC::DUX4 signature in the CDS PDX. The OS PDX did not express any OCTF signatures (**Fig. 2B**).

**Figure 2:**
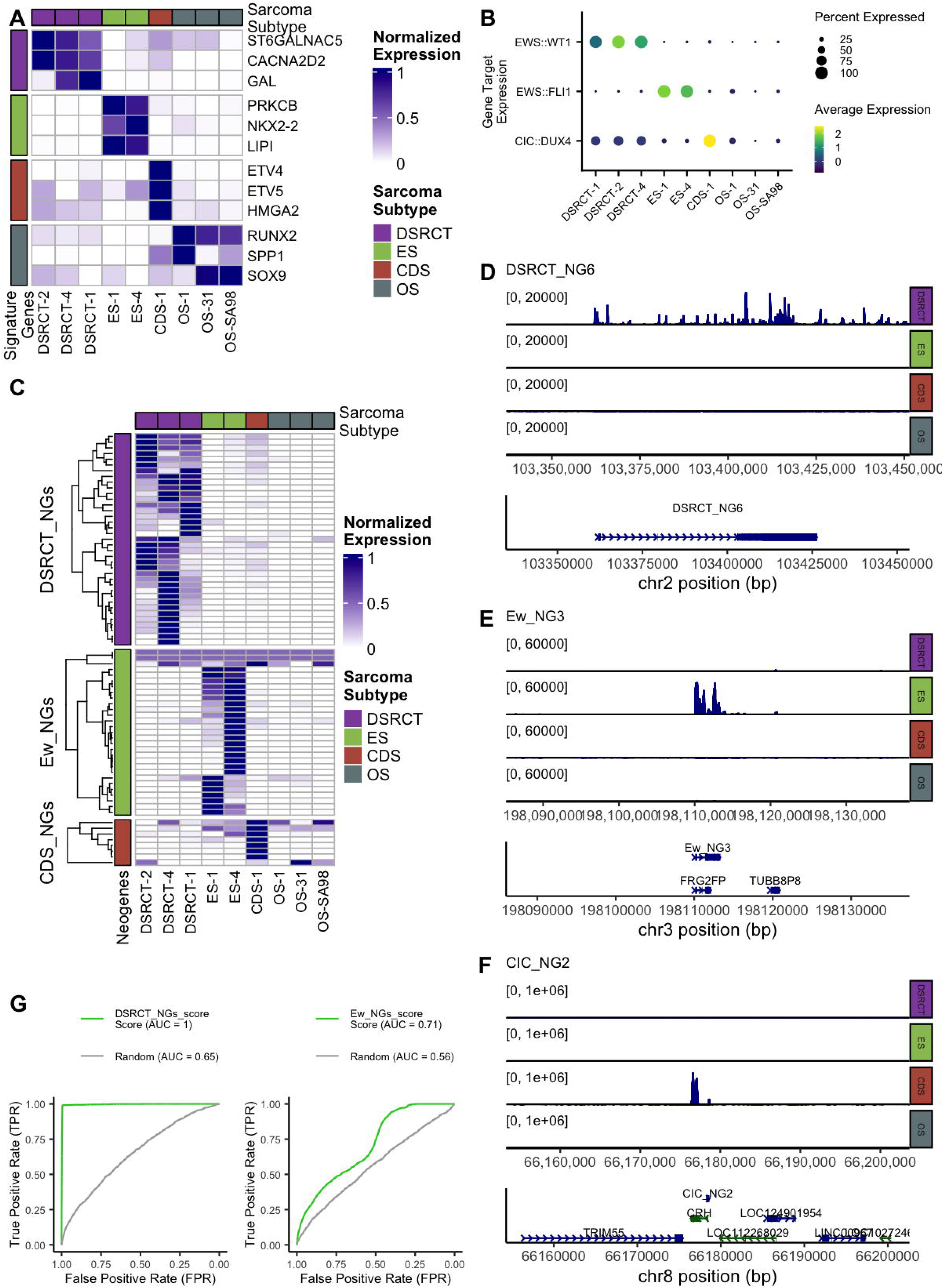
Selective and specific expression of neogenes in sarcoma subtypes. **A)** Heatmap comparing normalized gene expression of sarcoma-subtype-specific genes within DSRCT, ES, CIC::DUX4 sarcoma (CDS), and osteosarcoma (OS) PDX pseudo-bulk single-cell data. **B)** Dotplot comparing target gene expression of FPs EWS::WT1, EWS::FLI1, and CIC::DUX4. **C)** Heatmap comparing NG expression among sarcoma subtypes, demonstrating specific expression of NGs according to sarcoma subtype. **D)** Summary of ATAC-seq enrichment peaks of DSRCT_NG6, demonstrating DSRCT specificity. Horizontal tracks indicate enrichment in each PDX subtype, with subtype label on right. The lowest track shows the genomic location of NG and nearby gene loci. **E)** ATAC-seq enrichment of Ew_NG3, with a similar layout to panel D, demonstrating ES specificity. **F)** ATAC-seq enrichment of CIC_NG2, with a similar layout to panel D, demonstrating CDS specificity. **G)** Receiver operator characteristic curves for classification of DSRCT PDX tumor cells using DSRCT_NGs, Ew_NGs, and random genes, demonstrating high sensitivity and specificity only for DSRCT_NGs.

Next, we evaluated NG expression across our patient-derived xenograft (PDX) cohort. As anticipated, NGs specific to each OCTF were exclusively expressed in their corresponding sarcoma subtypes (**Fig. 2C**). In contrast, OS PDXs did not consistently express any NGs, reinforcing the specificity of NGs to OCTF-driven sarcomas. We observed inter-sample heterogeneity in NG expression within different DSRCT and ES PDXs, suggesting patient-specific variation in NG activation. Despite this variability, the data strongly support the subtype specificity of NGs. For example, DSRCT_NG6 expression was restricted to DSRCT PDXs, while Ew_NG3 was exclusively detected in ES PDXs, and CIC_NG2 was detected solely in CDS PDXs (**Fig. 2D, E, F**). To assess the specificity potential of NGs, we applied logistic regression models to classify cells from OCTF-driven PDXs based on NG expression (**Fig. 2G**). The dataset was randomly split into training and test sets to classify each sarcoma subtype. Using DSRCT_NGs, the model accurately identified DSRCT cells (AUC = 1.0), outperforming models based on randomly selected genes (AUC = 0.65) or Ew_NGs (AUC = 0.71). We further evaluated the prediction of ES from Ew_NGs and found similar performance (AUC = 0.94, **Supplementary** Fig. 3C). This was not observed for OS cells which had similar performances between Ew_NGs (AUC = 0.72) and randomly selected genes (AUC = 0.66) or DSRCT_NGs (AUC = 0.71) and randomly chosen genes (AUC = 0.67, **Supplementary** Fig. 3D). These findings highlight the potential of NGs as particular molecular classifiers for OCTF-driven sarcomas.

### Neogene expression is spatially enriched in tumor cells in DSRCT

To validate the spatial distribution of DSRCT_NGs, we performed RNAscope assays on FFPE tumor specimens from two DSRCT patients using the Lunaphore COMET™ Multiomics platform. Tumor cells were identified through marked expression of CACNA2D2, an established target of EWS::WT1^44^. Representative images of DSRCT_NG21 and DSRCT_NG33 showed puncta predominantly localized within tumor regions, as defined by threshold-based segmentation (**Fig. 3A**). Puncta were readily detectable at subcellular resolution, and higher magnification insets confirmed clear signals in tumor nests, which were also confirmed with DSRCT_NG6 and DSRCT_NG30 (**Supplementary** Fig. 4A), as well as the positive control probe UBC (**Supplementary** Fig. 4B). Quantification across all four DSRCT_NGs revealed consistently higher puncta density in the tumor region compared with stroma (**Fig. 3B**). Although absolute densities varied among probes, the increased enrichment was uniform, with each probe showing a several-fold increase in tumor regions. When benchmarked against positive control probes, DSRCT_NG puncta density was comparable to controls within tumor regions, indicating robust expression (**Fig. 3C**). In contrast, DSRCT_NGs exhibited a marked depletion in stroma, where puncta density was significantly reduced compared with positive controls (****, *P* = 1.20 × 10⁻L). By comparison, positive control probes maintained relatively stable densities across tumor and stroma, consistent with their role as broadly expressed housekeeping genes (ns, *P* = 0.135). Fold-change analysis further confirmed tumor enrichment, with DSRCT_NGs showing logL increases of ∼3–4. In contrast, positive control probes showed only modest enrichment or no difference (**Fig. 3D**). Together, these findings establish that DSRCT_NGs are spatially enriched within tumor cells, in contrast to the ubiquitous distribution of positive control transcripts, highlighting their tumor-cell specificity at the tissue level.

**Figure 3:**
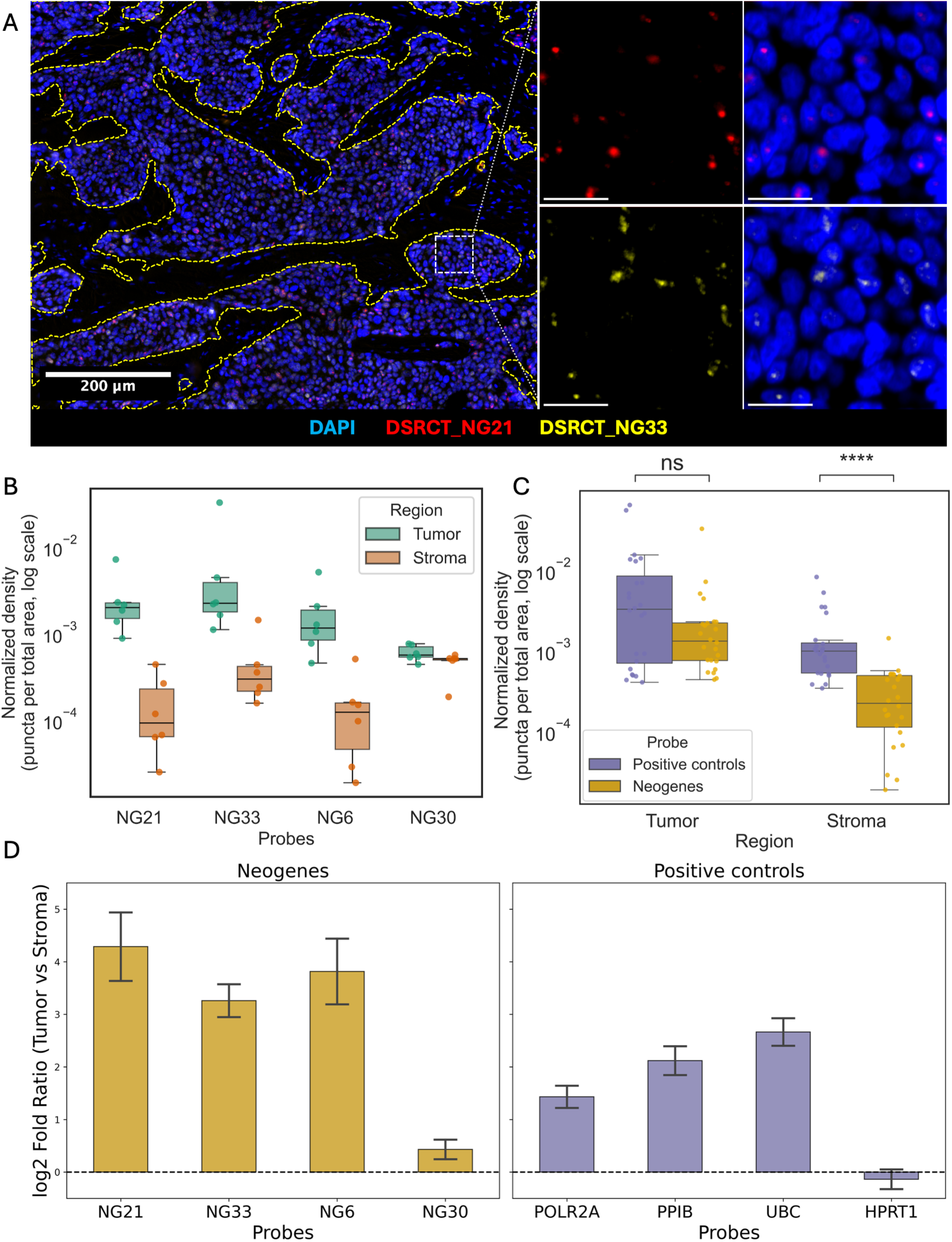
Tumor-cell-specific spatial expression of DSRCT neogenes. **A)** Representative image of RNAscope fluorescent in situ hybridization of DSRCT_NG21 (red) and DSRCT_NG33 (yellow) with DAPI (blue). A yellow segmentation border delineates tumor and stroma. Insets show higher magnification. Scale bars: 200 µm (left), 20 µm (right). **B)** Boxplots of normalized puncta density (puncta/µm², log scale) for four DSRCT_NGs (NG21, NG33, NG6, NG30) in tumor versus stroma regions. **C)** Comparison of puncta density between neogenes and positive control probes, showing significantly lower density for neogenes in stroma (****, *P* = 1.20 × 10LL) but not in tumor (ns, *P* = 0.135; Mann-Whitney U test). **D)** LogL fold ratio (tumor vs. stroma) of puncta density for neogenes and positive controls, with most probes enriched in tumor except for HPRT1.

### Neogene expression is differentially driven by EWS::WT1 isoforms

Having established the specificity of DSRCT_NGs, we sought to characterize their regulation in response to EWS::WT1 FP knockdown (KD). We recently established and performed RNA-seq on a cohort of four DSRCT cell lines (JN-DSRCT-1, BER-DSRCT, SK-DSRCT2, BOD-DSRCT) that deplete FP utilizing a doxycycline(dox)-inducible shRNA system^15^. Analyzing NG expression within this RNA-seq data set, we found that KD of the EWS::WT1 FP led to a consistent reduction in DSRCT-specific NGs (**Fig. 4A**). While there was variability in EWS::WT1 FP KD sensitivity, we identified 13/37 NGs that decreased by at least ≥50% (log2FC ≤ -1) in all four cell lines and an additional 8/37 NGs with ≥50% reduction across three cell lines (**Fig. 4B**). The BOD-DSRCT and SK-DSRCT2 cell lines exhibited the largest number of NGs reduced after KD, with 29/37 and 25/37 NGs having ≥50% expression reduction, respectively.

**Figure 4:**
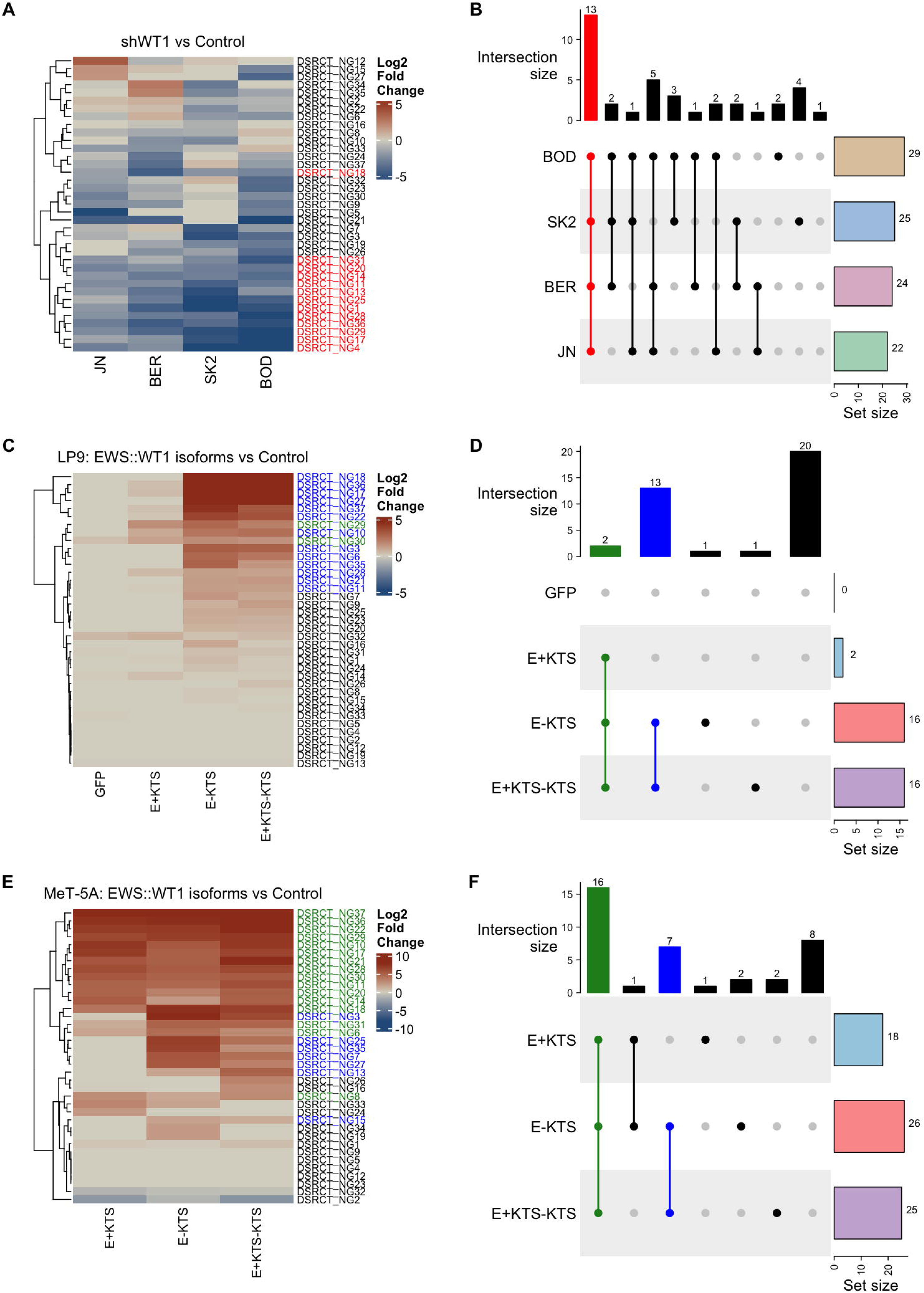
EWS::WT1 isoforms exhibit variable control of DSRCT neogene expression. **A)** Heatmap of log_2_-fold change in normalized gene expression in DSRCT_NGs caused by knockdown of EWS::WT1 (shWT1) relative to control shRNA in four DSRCT cell lines (JN-DSRCT-1, BER-DSRCT, SK-DSRCT2, BOD-DSRCT; shown abbreviated; each with n=2 replicates). **B)** UpSet plot showing compound overlap of differentially expressed neogenes (≥50% decrease) across four DSRCT cell lines in response to shWT1 knockdown relative to control shRNA. Neogenes affected in all four cell lines are also displayed in red in panel A. **C)** Heatmap of log_2_-fold change in normalized gene expression in DSRCT_NGs in LP9 cells transduced with GFP, E+KTS, E–KTS, or both isoforms, relative to nontransduced control cells (*n*=2 replicates each). **D)** UpSet plot showing compound overlap of differentially expressed neogenes across LP9 conditions relative to control shRNA. Neoenes affected in all isoform conditions or only E–KTS conditions are also displayed in green and blue, respectively, in panel C. **E)** Heatmap of log_2_-fold change in normalized gene expression in DSRCT_NGs in MeT-5A cells transduced with E+KTS, E–KTS, or both isoforms, relative to nontransduced control cells (*n*=3 replicates each). **F)** UpSet plot showing compound overlap of differentially expressed neogenes across MeT-5A conditions relative to control shRNA. Neogenes affected in all isoform conditions or only E–KTS conditions are also displayed in green and blue, respectively, in panel E.

The EWS::WT1 fusion protein (FP) exists in two isoforms, E+KTS and E–KTS, distinguished by alternative splicing within the WT1 domain^45^; this splicing event includes or excludes three amino acids (lysine, threonine, and serine; KTS) between zinc fingers 3 and 4. In another recent study, we transduced one or both isoforms into the LP9 mesothelial cell line. The E–KTS isoform is more transcriptionally active, driving most EWS::WT1-regulated gene expression changes^15^. Similarly, Boulay et al. performed isoform-specific transductions in the p53-mutant mesothelial cell line MeT-5A^46^. Using these two datasets, we assessed the contribution of each isoform to DSRCT_NG expression. In transfected LP9 cells, only 2 out of 37 DSRCT_NGs were upregulated (log2FC > 1) by E+KTS, while 16 out of 37 were induced by E–KTS; the same 16 genes were also induced when both isoforms were co-transduced (**Fig. 4C, D**). Notably, the two genes activated by E+KTS were also induced by E–KTS. In MeT-5A cells, E+KTS induced 18 of the 37 DSRCT_NGs, E–KTS induced 26, and co-expression of both isoforms induced 25 (**Fig. 4E, F**). All 16 NGs activated by E–KTS in LP9 cells were also upregulated by E–KTS in MeT-5A. Among the 10 additional NGs induced by E–KTS in MeT-5A, five were also responsive to E+KTS, while the remaining five were uniquely induced by E–KTS. These findings reveal a consistent set of DSRCT_NGs activated by the E–KTS isoform across both cell lines, highlighting its dominant role in driving DSRCT_NG expression compared to E+KTS.

### Neogene motif enrichment highlights EWS::WT1 isoform-dependent epigenetic regulation

To gain deeper insight into the role of EWS::WT1 in regulating DSRCT_NGs, we conducted HOMER motif enrichment analysis on regions spanning ±1000 bp from the transcription start site (TSS) of each NG. Supporting a direct regulatory function of EWS::WT1, the WT1 consensus binding motif, CTCCC(A/C)C, emerged as the most significantly enriched motif within these regions (**Fig. 5A**, **Supplementary Table 3**). Using HOMER to evaluate de novo motifs, we also observed TCC motif repeats enriched in 46.7% of the DSRCT_NG as described before^46^. To test the specific FP binding of the DSRCT_NGs, we evaluated two independent sets of ChIP-seq data^46,47^ examining the binding of EWS::WT1 in the JN-DSRCT-1 cell line (**Fig. 5B**). We found that 58.9 % of DSRCT_NGs have an EWS::WT1 binding site within 1k bps of the TSS (**Fig. 5C**). In contrast, only 3.1% of non-DSRCT_NGs regulated by other OCTFs had an EWS::WT1 binding site within 1 kb of the TSS. Analysis of key histone modifications revealed that H3K4me1, H3K4me3, H3K9ac, and H3K27ac were present near the TSS in approximately 75% of DSRCT NGs. In contrast, the repressive mark H3K27me3 was detected in only 2% of these NGs.

**Figure 5:**
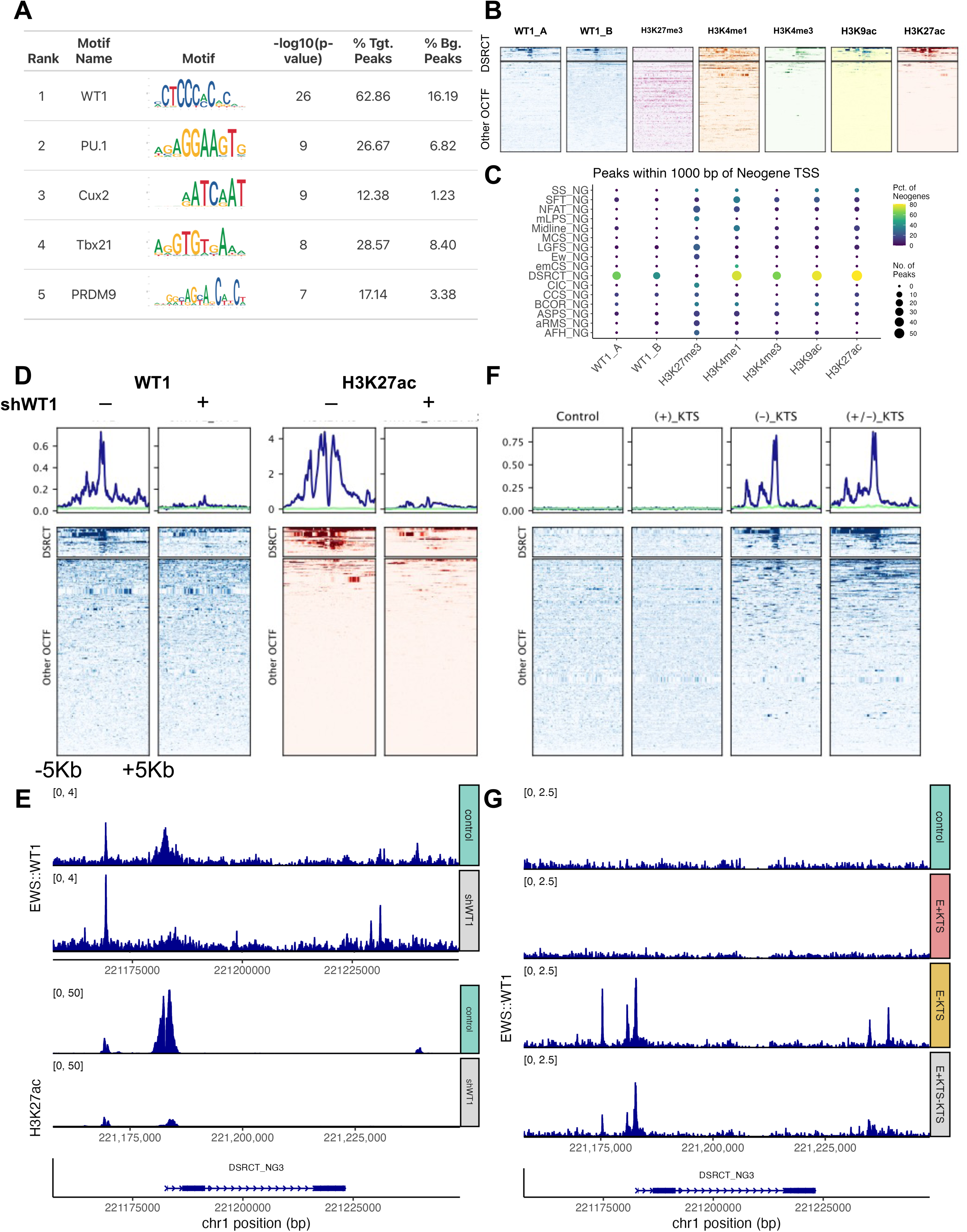
EWS::WT1 binding controls DSRCT neogene expression. **A)** Table of top enriched transcription factor motifs for DSRCT_NG transcription start site (TSS), including motif name, motif sequence, enrichment p-value, percent target and background peaks. **B)** Heatmap visualization of WT1 and histone marks at DSRCT_NG loci and other OCTF NG loci. **C)** Dotplot of enriched peaks among sarcoma-subtype-specific NG, with color denoting percent of NGs expressed and dot size denoting number of peaks enriched within 1000 base-pairs of NG TSS. **D)** Enrichment of EWS::WT1 binding at DSCRT NGs and other NG loci in control and after WT1 knockdown. **E)** Enrichment of H3K27Ac histone marker at DSCRT NG loci and other NG loci in control and after WT1 knockdown. **F)** Enrichment of EWS::WT1-HA after HA pull down for different EWS::WT1 KTS isoforms at DSCRT NG and other NG loci. **G)** EWS1::WT1 and H3K27Ac ChIP-seq coverage tracks about DSRCT_NG 3, before and after WT1 knockdown. G) Coverage tracks of EWS::WT1 KTS isoforms at DSRCT_NG 3.

Next, we compared ChIP-seq profiles of WT1 binding and H3K27Ac marker in the JN-DSRCT-1 cell line, comparing control samples to those treated with shRNA targeting the EWS::WT1 fusion protein (**Fig. 5DE**). Treatment with the shRNA led to an approximately fourfold reduction in the number of peaks associated with the TSS of DSRCT_NGs (**Supplementary** Fig. 5A). We also confirm a strong enrichment of WT1 binding near the TSS of DSRCT_NGs as previously described^1^. Additionally, we observed enhanced binding of the active histone marker H3K27ac at DSRCT_NGs compared to NGs from other OCTF-driven cancer subtypes. Interestingly, upon KD of EWS::WT1, we observed decreased enrichment of H3K27ac, which suggests that loss of the FP reduced chromatin accessibility at enhancer regions for the DSRCT_NG loci.

Similarly, transduction of specific EWS::WT1 isoforms to the MeT-5A cell line variably enriched peaks near the TSS of DSRCT_NGs. We found that 62.2% of DSRCT_NGs have an EWS::WT1 binding site within 1k bps of the TSS for E–KTS (**Fig. 5FG** and **Supplementary** Fig. 5B). In contrast, only 41.9% of DSRCT_NGs were detected in E+KTS. Unsurprisingly, the E+/–KTS condition was also enriched for 62.2% of DSRCT_NGs peaks near the TSS, suggesting again that the E–KTS dominantly drives DSRCT_NG expression. Interestingly, when we evaluated wildtype WT1 with +KTS or –KTS isoforms (lacking the EWS fusion), we observed a marked decrease in DSRCT_NG peaks (WT1–KTS = 20.3% and WT1+KTS = 0%, **Supplementary** Fig. 5C). This was also evident in the lack of peaks in the coverage plots (**Supplementary** Fig. 5DE).

### Single-cell multiomics reveals differential chromatin accessibility in DSRCT cell line

Next, we analyzed single-cell multiome data from the JN-DSRCT-1 and TC71 (ES) cell lines to investigate chromatin accessibility patterns that may regulate NG expression. This integrative approach allowed us to simultaneously assess gene expression and chromatin accessibility at the single-cell level. We confirmed the expected expression profiles characteristic of DSRCT and ES, including the expression of OCTF-specific NGs in their respective cell lines (**Fig. 6A**).

**Figure 6:**
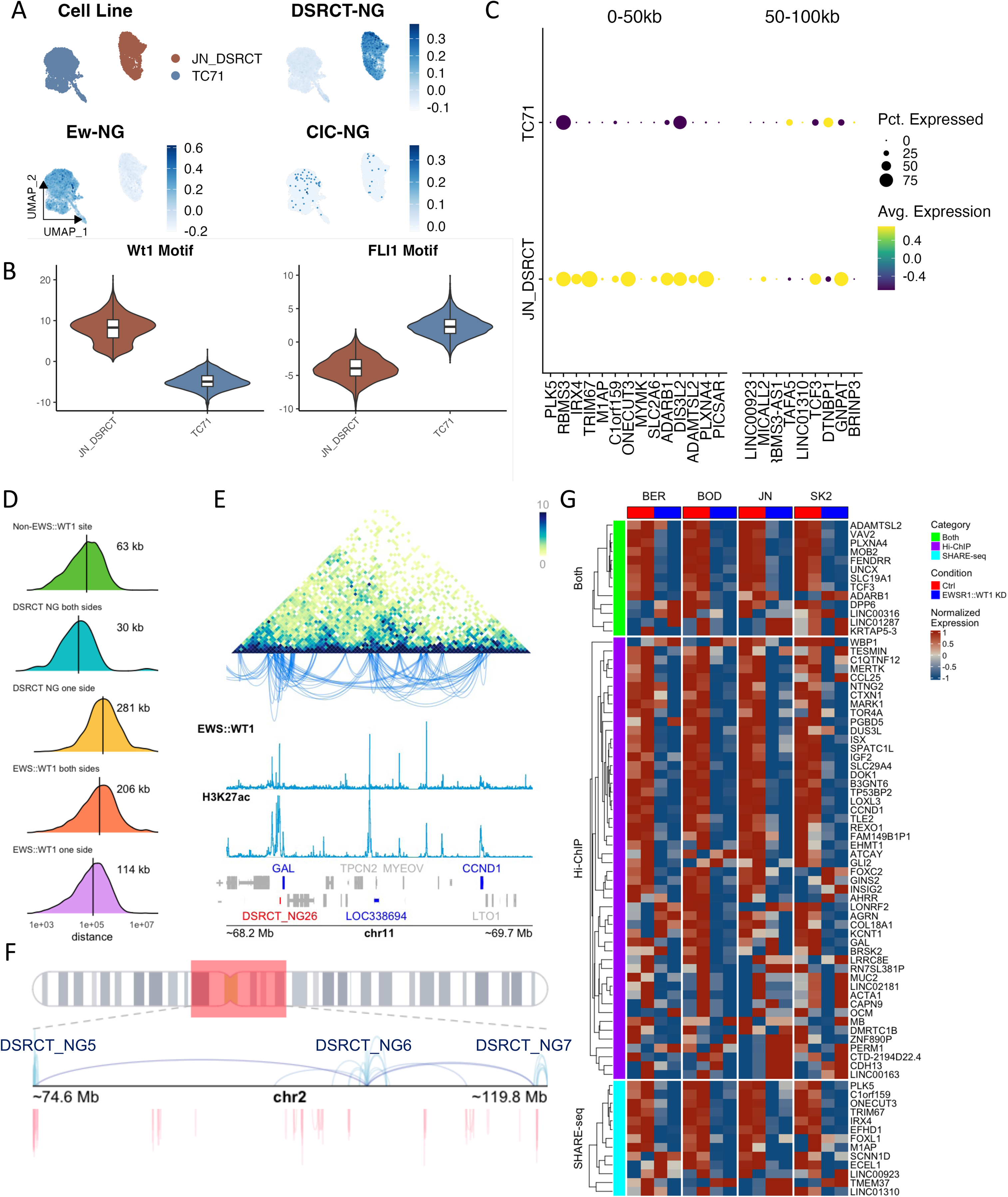
Enhancer-promoter hubs at DSRCT neogene loci control nearby gene targets. **A)** UMAP projection for JN-DSRCT-1 and TC71 (ES) cell lines. Subpanels demonstrate specific DSRCT and ES NGs in respective cell lines and no expression of CIC::DUX4 NGs. **B)** Violin plots of chromVAR-inferred WT1 and FLI1 motif expression in JN-DSCRT-1 and TC71 cell lines. **C)** Dotplot of gene located within 0-50 kilobases and 50-100 kilobases of DSRCT_NGs in JN-DSRCT and TC71 cell lines. **D)** Ridgeline plot of looping distance at anchors within H3K27ac Hi-ChIP data relative to Non-EWS::WT1 sites, DSRCT_NG sites, and EWS::WT1 loci. **E)** Chromatin interaction matrix near the DSRCT_NG26 locus in chromosome 11 with loops to nearby DSRCT genes, like GAL and CCND1. Tracks indicate EWS::WT1 and H3K27Ac peaks along with gene annotations. **F)** Example of chromosome 2 map indicating long-range interactions originating from DSRCT_NGs (NG5, NG6, and NG7). Dark red indicates a non-EWS::WT1 site, and dark blue indicates DSRCT_NG involvement. **G)** Heatmap of normalized bulk gene expression of differentially expressed genes in response to WT1 knockdown relative to control shRNA. Genes near DSRCT_NG are organized relative to prior identification within Hi-ChIP, SHARE-seq, or both datasets. Expression of genes near DSRCT_NG is reduced upon WT1 knockdown.

To explore the regulatory landscape, we identified accessible chromatin regions and performed motif enrichment inference. Notably, JN-DSRCT-1 cells exhibited a higher enrichment of the WT1 transcription factor motif compared to TC71 cells, consistent with the presence of the EWS::WT1 fusion protein that defines DSRCT (**Fig. 6B**). This finding supports the role of WT1 in shaping the chromatin landscape in DSRCT. Furthermore, we evaluated chromatin accessibility in representative cell lines for NGs (**Supplementary** Fig. 6A). The DSRCT_NGs were accessible only in DSRCT cell lines. This observation was also shared with other NGs for respective OCTF (**Supplementary** Fig. 6B). This suggested that the FP may play a role in modifying chromatin accessibility at the NG loci.

Focusing on DSRCT_NGs, we observed substantial heterogeneity in their expression across individual cells. We hypothesized that this variability is driven by differential chromatin accessibility at the NG loci. To test this, we employed the SHARE-seq framework to infer peak-to-gene associations by correlating chromatin accessibility with gene expression across single cells. This analysis provided insights into the regulatory architecture underlying NG expression and highlighted potential cis-regulatory elements contributing to the observed transcriptional heterogeneity^48,49^. We identified 422 significant peak-gene associations within 50 kb around the TSSs for DSRCT_NGs (**Supplementary Table 4A**, p < 0.05). We identified that several peak-gene associations originated from the TSS of the DSRCT_NGs to nearby genes (**Supplementary** Fig. 7A). Interestingly, this was marked by both EWS::WT1 binding and H3K27ac in the JN-DSRCT-1 cell line. We investigated genes both within 50 kb and between 50 kb and 100 kb from the DSRCT_NG TSS, finding that these genes were strongly expressed in JN-DSRCT-1 compared to the ES cell line TC71 (**Fig. 6C, Supplementary Table 4B**).

To better understand the regulatory dynamics of EWS::WT1 near DSRCT_NG loci, we analyzed H3K27ac-mediated Hi-ChIP data in the JN-DSRCT-1 cell line. Boulay et al. showed that EWS::WT1-associated chromatin loops often span large genomic distances^46^. Using this data, we identified chromatin loops with EWS::WT1 binding sites and/or DSRCT_NG loci on either or both sides of the loop. Consistent with prior findings of long-range EWS::WT1 interactions, we observed that loops involving DSRCT_NGs also tend to cover long distances. Specifically, when DSRCT_NGs were present on only one side of a loop, the median distance was 281 kb, substantially greater than loops involving non-EWS::WT1 sites (63 kb) or EWS::WT1 sites on one side that do not involve DSRCT_NGs (114 kb) (**Fig. 6D, Supplementary Table 5A**). When DSRCT_NG loci were present on both sides of the loop, the interactions were typically short-range and often occurred within the same DSRCT_NG. However, in some cases, we observed long-range inter-DSRCT_NG looping within the identical chromosomes, forming pairs such as DSRCT_NG6/7, DSRCT_NG13/14, DSRCT_NG23/24, DSRCT_NG28/29, and DSRCT_NG30/31. Notably, loops that included DSRCT_NGs on one side or EWS::WT1 binding sites on both ends (206 kb) had long-range interactions (**Fig. 6D**). Notably, the DSRCT_NG promoter regions were involved in significantly more looping interactions than typical EWS::WT1 gene targets, with median loop counts of 10 and 0, respectively (**Supplementary Table 5BC**). This led to our observation that some DSRCT_NGs are linked to DSRCT marker genes, like *GAL* and *CCND1.* This can be seen in the chromatin interaction matrix showing the DSRCT_NG26 locus on chromosome 11, where the EWS::WT1 ChIP signal is at the TSS of DSRCT_NG26 and H3K27ac is at *GAL* and *CCND1* loci (**Fig. 6E).** We further evaluated the genomic sites that are linked to DSRCT_NGs (**Fig. 6F and Supplementary** Fig. 7B).

To comprehensively identify genes associated with loops involving DSRCT_NGs, we used both SHARE-seq and Hi-ChIP. From this set, we identified genes that were significantly downregulated or upregulated in DSRCT cell lines after EWS::WT1 knockdown, with an absolute log2 fold change of ≥1.5 (**Fig. 6G**). These same genes are upregulated in the LP9 mesothelial cell lines expressing the E+KTS, E– KTS, or both isoforms (**Supplementary** Fig. 8A). However, like the prior data, it appears that the E–KTS is driving most of the gene expression. Importantly, most of these genes that are linked to the DSRCT_NGs are specific to DSRCT (**Supplementary** Fig. 8B). We evaluated the overlap of the promoter region of DSRCT_NGs and the linked genes with the EWS::WT1 binding sites or H3K27ac in JN-DSRCT-1 by a permutation test. We found that there was a stronger association, denoted by the Z-score, for DSRCT_NGs than for the linked genes to EWS::WT1, with nearly 57% of DSRCT_NGs found to overlap with EWS::WT1 binding compared to 12% for the linked genes, yet we know the linked genes are highly expressed in DSRCT (**Supplementary Table 5DE**). These data reinforce the finding that the EWS::WT1 fusion protein regulates some gene targets through a complex network of chromatin interactions involving DSRCT_NG loci, which harbor the EWS::WT1 binding site.

## Discussion

The recent discovery of disease-specific NGs by Vibert et al., who demonstrated a remarkable specificity of NGs to corresponding OCTF fusion proteins, has sparked significant interest in exploring their biological roles, regulatory mechanisms, and therapeutic potential. While initial investigations have primarily focused on the function of NGs in ES, our research has extended this line of inquiry to DSRCT. In doing so, we have begun to elucidate the epigenetic regulation of DSRCT_NGs, uncovering novel insights into their expression patterns and potential as biomarkers or therapeutic targets. This expanded focus broadens our understanding of NG biology across fusion-driven sarcomas and highlights their emerging relevance in clinical and translational research contexts.

Our evaluation of NGs in patients, PDX, and cell line models extends their specificity for OCTF in a clinically and translationally relevant context. In patient samples, DSRCT_NGs were restricted to specimens with EWS::WT1 FP; in one patient with multiple tumors sampled, only those with detectable FP expressed DSRCT_NGs. Analysis of PDXs also revealed that NGs are selectively expressed in OCTF-driven sarcomas, with distinct NG profiles corresponding to EWS::WT1, EWS::FLI1, and CIC::DUX4 fusion proteins. In fact, these NG sets alone were sufficient to construct molecular classifiers of their respective disease subtypes. Additionally, spatial staining of NG puncta also indicated specific expression of DSRCT NGs in tumor cells and not in stroma. Both modalities show we can build transcriptomic and imaging classifiers for tumor cells.

Given that the NGs are purported to be regulated by OCTF, we investigated the regulation of DSRCT_NGs to better elucidate the epigenetic consequence on DSRCT biology by the EWS::WT1 FP. Knockdown of the EWS::WT1 fusion protein consistently reduced expression of DSRCT_NGs across four cell lines, confirming their regulation by the fusion protein. Further isoform-specific analysis revealed that the E–KTS isoform dominantly drives DSRCT_NG expression across multiple cell lines, underscoring its critical role in DSRCT transcriptional programming. This isoform specificity was not only observed in expression data, but also in examination of the binding of the two isoforms, wherein the E–KTS isoform bound strongly to NG promoter regions while the E+KTS isoform was absent.

Further multiomic analysis revealed short- and long-range genomic interactions, underscoring a complex epigenetic landscape governing NG expression in DSRCT. We uncovered a distinctive chromatin architecture characterized by extensive looping interactions between DSRCT_NG loci and nearby genes, expanding upon prior observations of EWS::WT1-mediated chromatin looping. Identifying peak-to-gene associations and enhancer-promoter hubs linked to NG expression provides mechanistic insight into how EWS::WT1 orchestrates the DSRCT transcriptome^50,51^. Strikingly, WT1 motif enrichment was confined to NG loci. In contrast, nearby gene loci exhibited strong H3K27ac signals, suggesting that EWS::WT1 may act as a pioneer factor, analogous to EWS::FLI1 in Ewing sarcoma, by remodeling chromatin to activate lineage-inappropriate transcriptional programs. These findings support a broader paradigm in which EWS fusion proteins hijack chromatin topology to drive oncogenic gene expression. Notably, the specificity of NG-linked chromatin loops to DSRCT, coupled with their responsiveness to EWS::WT1 knockdown, highlights their potential as biomarkers for disease stratification and as targets for therapeutic intervention.

Immunotherapy has transformed the treatment of many solid cancers, especially those with high tumor mutational burden, which is a predictor of response to immune checkpoint inhibitors. However, sarcomas are generally not great candidates for immunotherapies, except for perhaps alveolar soft part sarcoma, undifferentiated pleomorphic sarcoma, and dedifferentiated liposarcoma^52^. DSRCT and many other sarcomas have a single gene translocation and low tumor mutational burden, which is predictive of low response to immune checkpoint inhibitors. Work from our group investigated the immunogenic profile of DSRCT, showing that DSRCT belonged to the immunologically ‘cold’ group^13^. Like other types of immunologically cold tumors, eliciting an immune response in DSRCT may require stimulation by exogenous factors such as a cancer vaccine, CAR T-cell, or NK cell-based therapies^53^.

Recent findings from our group and others have reignited interest in exploring immunotherapeutic strategies for DSRCT. The Delattre group demonstrated that certain Ew_NGs generate ES-specific peptides presented by MHC Class I molecules on ES cells^54^. Importantly, cytotoxic T cells targeting these peptides selectively killed ES cells, but not those with EWS::FLI1 knockdown, indicating that peptide presentation depends on the fusion oncogene. Complementing this, Banks et al. identified a neoantigen derived from the EWS::WT1 fusion junction in DSRCT, leading to the characterization of a 9-amino acid peptide spanning the fusion site recognized by CD8L T cells^55^. Although DSRCT-specific peptides derived from DSRCT_NGs have yet to be fully characterized, these studies underscore the feasibility of targeting fusion-driven neoantigens in DSRCT.

By integrating single-cell multiomics with bulk RNA-seq, patient-derived xenograft (PDX) models, and primary patient tumor data, this study provides a comprehensive view of the transcriptional and epigenetic landscape of DSRCT. Through chromatin conformation analyses including Hi-ChIP and ChIP-seq, we reveal how the EWS::WT1 fusion protein orchestrates long-range enhancer-promoter interactions and selectively remodels chromatin accessibility at NG loci. Knockdown experiments further validate the functional relevance of these regulatory circuits, demonstrating that EWS::WT1 directly influences the expression of both NG and their linked gene targets. This integrative framework advances our understanding of OCTF by uncovering the multi-layered mechanisms through which EWS::WT1 hijacks chromatin topology to activate oncogenic programs. The methodologies and insights presented here, spanning patient samples, in vivo models, and multimodal sequencing, are broadly applicable to other fusion-positive malignancies and lay the groundwork for precision epigenetic therapies targeting aberrant chromatin architecture.

## Acknowledgements

The authors acknowledge the High Performance Computing for Research facility at The University of Texas MD Anderson Cancer Center for providing the computational resources that supported the research reported in this paper. We also thank the Institutional Tissue Bank and the Cancer Prevention Research Institute of Texas Pediatric Solid Tumors Comprehensive Data Resource Core (RP180819) for providing tissue specimens and Energy Transfer Partners for their additional support in collecting and storing these specimens.

