## Supplemental Material for "EWS::WT1 Isoform-Dependent Regulation of Neogenes in Desmoplastic Small Round Cell Tumors"

Supplemental Figures

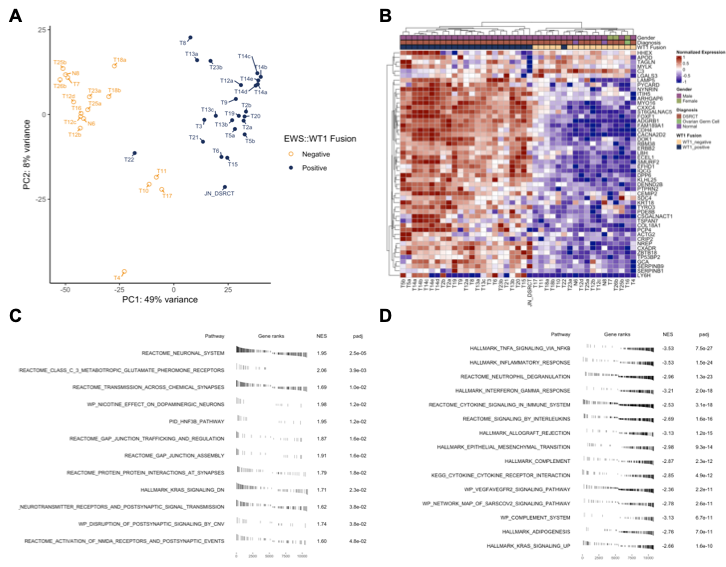

**Supplementary Figure 1. Transcriptomic differences in DSRCT specimens with detectable EWS::WT1 fusion.** A) Principal component analysis (PCA) of 40 DSRCT specimens reveals clustering based on fusion status. Most fusion-positive specimens cluster together, while fusion-negative and ambiguous specimens (e.g., T22) show distinct separation. B) Differential gene expression analysis highlights fusion-specific genes such as *ST6GALNAC5*, *CACNA2D2*, *GAL*, and *GALP* enriched in fusion-positive specimens. C) Gene set enrichment analysis (GSEA) identifies neuronal and synaptic pathways enriched in fusion-positive specimens. D) Pathways related to immune response, inflammation, and adipogenesis are downregulated in fusion-positive specimens.

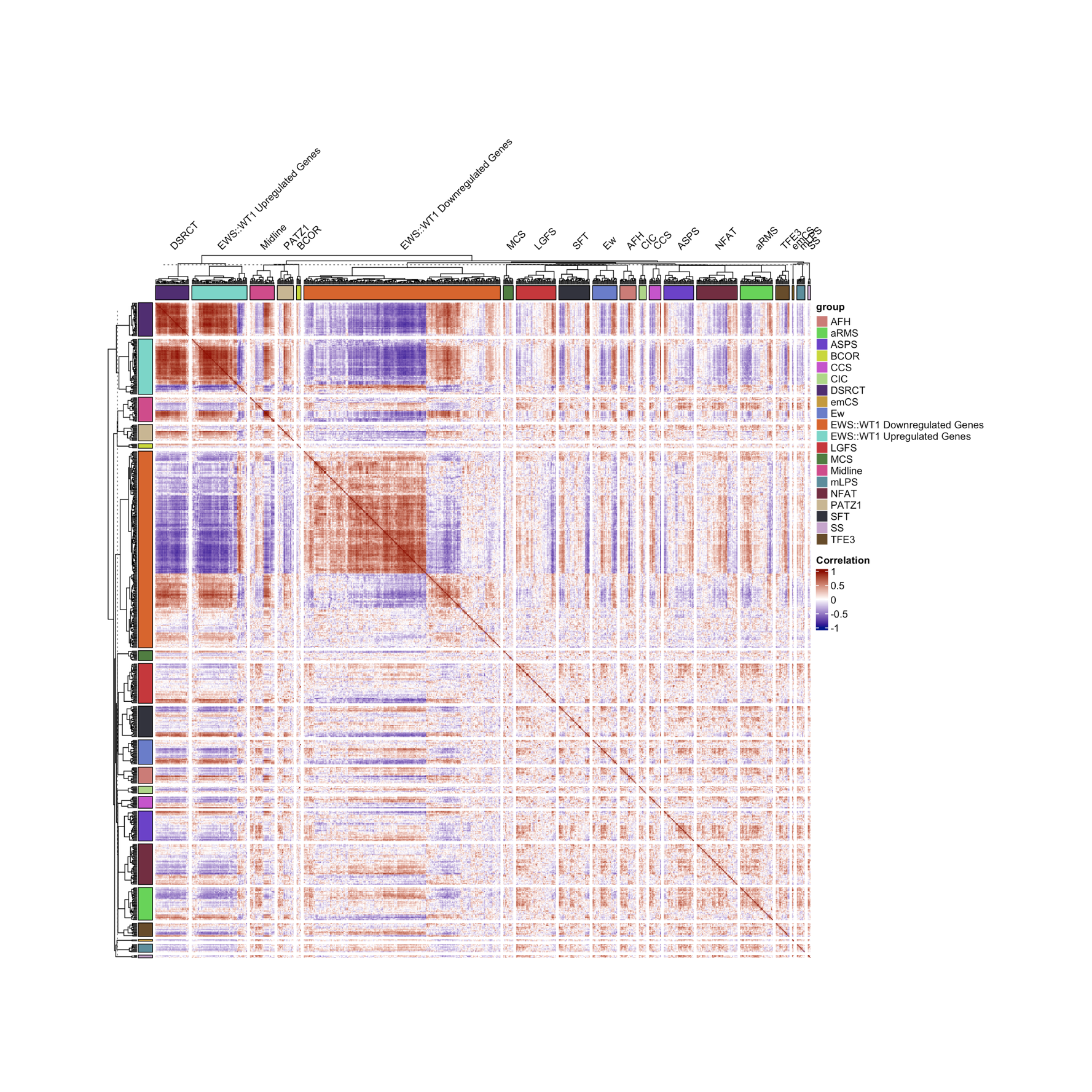

**Supplementary Figure 2. Correlation between DSRCT_NGs and EWS::WT1-regulated genes.** Gene-gene correlation analysis shows a strong positive correlation between DSRCT_NGs and genes upregulated by the EWS::WT1 fusion protein. Conversely, genes downregulated by EWS::WT1 exhibit a negative correlation with DSRCT_NGs.

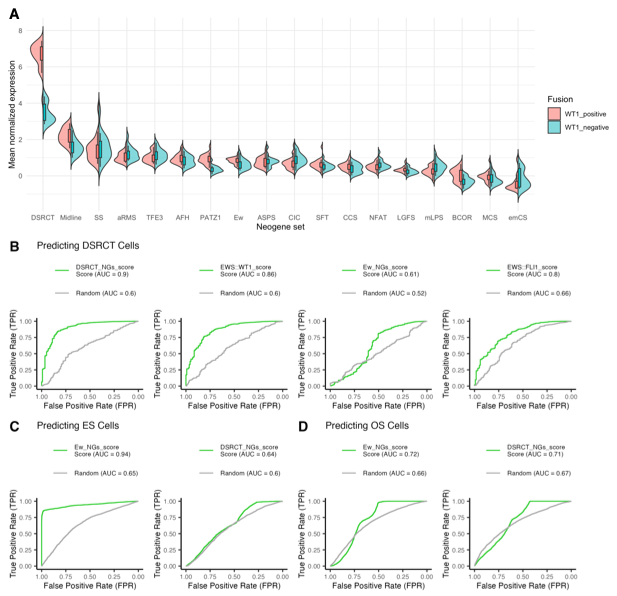

**Supplementary Figure 3. Specificity of DSRCT_NGs compared to other OCTF-driven neogenes.** **A)** Expression analysis of neogenes regulated by other oncogenic transcription factors (OCTFs) demonstrates that DSRCT_NGs are uniquely and consistently expressed in EWS::WT1 fusion-positive specimens. In contrast, other NGs show weaker expression. Split violins demonstrate distribution of average disease-specific neogene expression within fusion-positive or fusion-negative specimens. **B)** Receiver Operating Characteristic (ROC) curve analyses performed to classify DSRCT cells in single-nucleus RNA-seq data. Signatures for DSRCT_NGs, EWS::WT1, ES_NGs, and EWS::FLI1 were calculated in Seurat using AddModuleScore. The results show that DSRCT_NGs can reliably predict DSRCT cells unlike Ew_NGs or EWS::FLI1 score. **C)** Similar performance was found for ES PDX cells using Ew_NGs. **D)** OS cells could not be reliably predicted using either Ew_NGs or DSRCT_NGs.

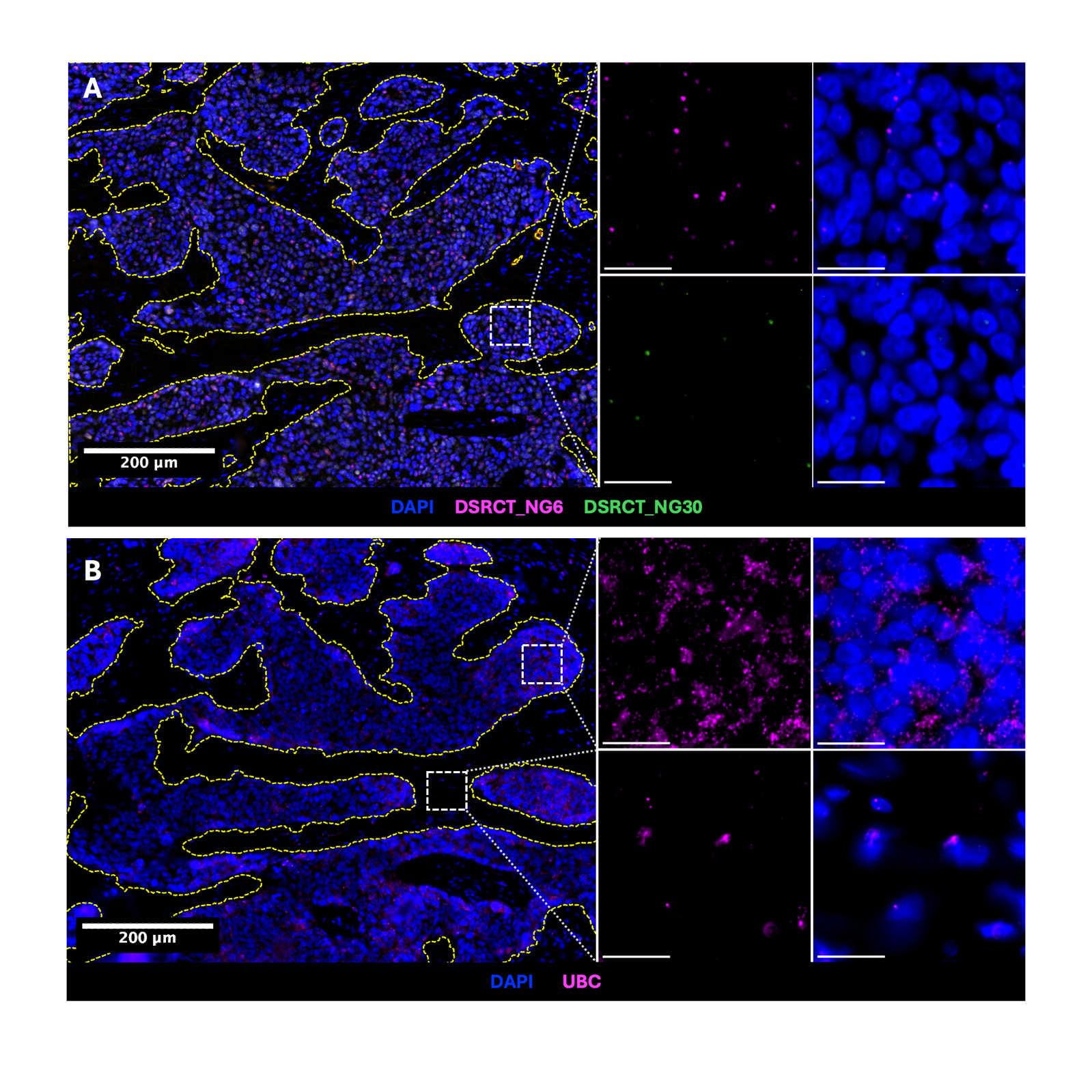

**Supplementary Figure 4. A)** Representative RNAScope fluorescent in situ hybridization image of DSRCT_NG6 (magenta) and DSRCT_NG30 (green) with DAPI (blue). **B)** Representative RNAScope fluorescent in situ hybridization image of positive control probe UBC (magenta) with DAPI (blue). Tumor and stroma are delineated by a yellow segmentation mask. Insets show higher magnification. Scale bars: 200 µm (left), 20 µm (right).

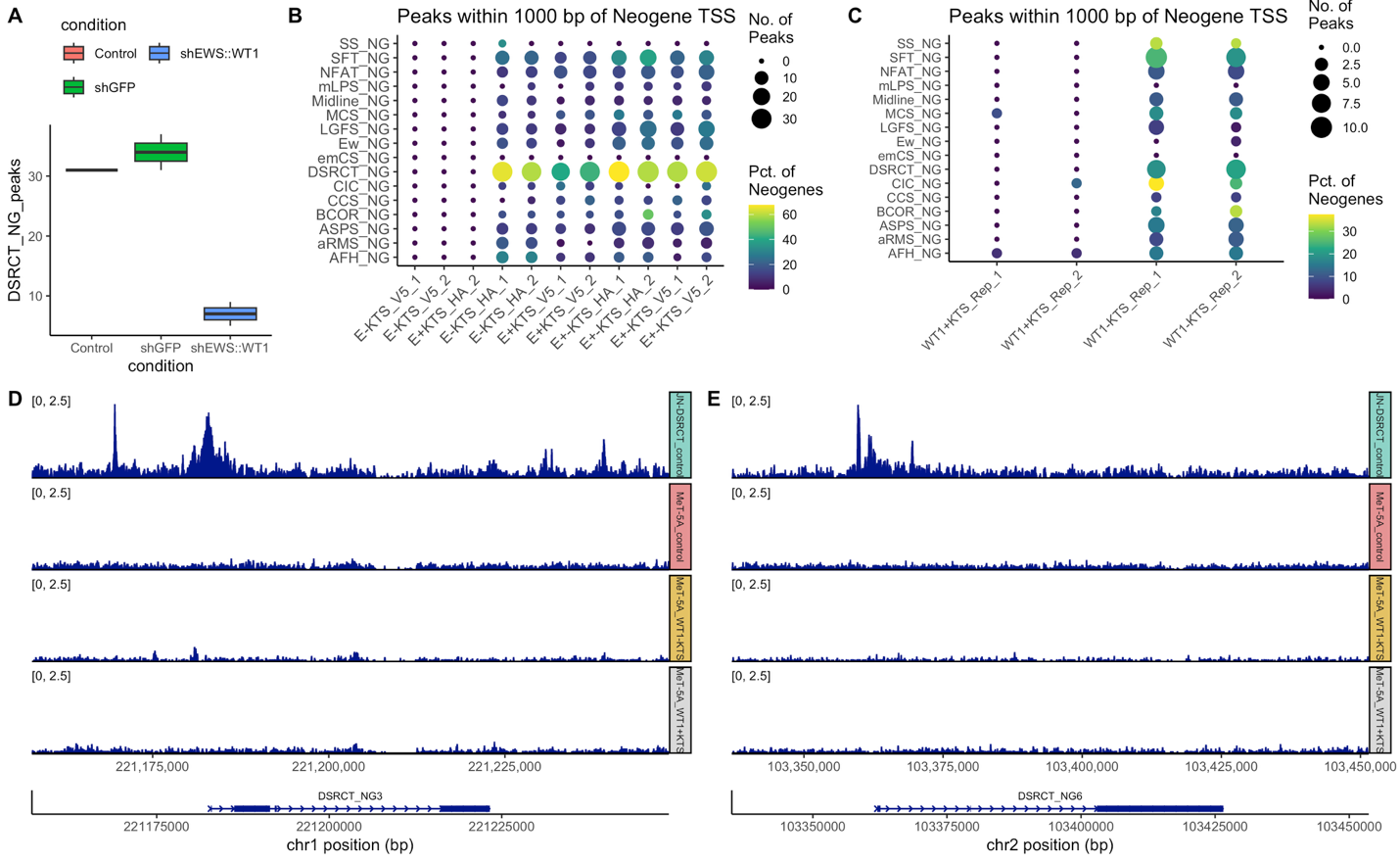

**Supplementary Figure 5. WT1 isoform lacks binding affinity of EWS::WT1 isoforms.** **A)** Drop in DSRCT_NG peaks near WT1 binding site upon EWS::WT1 knockdown. **B)** Percentage of DSRCT_NGs with EWS::WT1 binding sites within 1 kb of the TSS across shRNA knockdown conditions. **B)** Isoform-specific EWS::WT1 binding peaks within 1000 bp of neogene TSS across several fusion-driven cancer subtype neogene sets. E–KTS conditions show the highest enrichment. **C)** Isoform-specific WT1 binding peaks within 1000 bp of neogene TSS across several fusion-driven cancer subtype neogene sets. WT1–KTS shows markedly increased binding relative to WT1+KTS. **D)** Comparison of WT1 isoform binding near DSRCT_NG3 TSS in WT1 isoform-transduced Met5a cells and JN-DSRCT-1. WT1–KTS shows limited binding; WT1+KTS shows virtually none. **E)** Similar to panel D, coverage plots for DSRCT_NG6 showing lack of WT1 isoform binding.

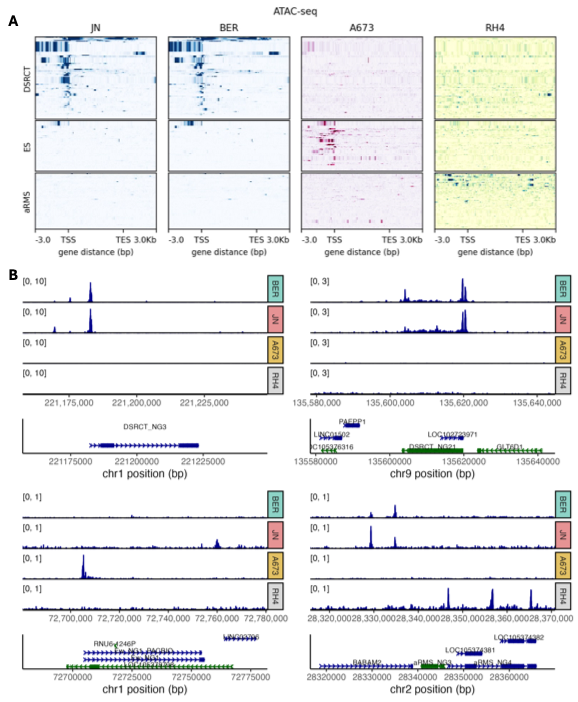

**Supplementary Figure 6. Chromatin accessibility of NGs in different sarcoma subtypes. A)** Heatmap of accessible peaks for DSRCT, ES, and aRMS NGs for respective cell lines of DSRCT (JN, BER), ES (A673), and aRMS (RH4) subtypes. **B)** Representative NG loci (DSRCT_NG3, DSRCT_NG21, Ew_NG1, aRMS_NG3) and ATAC-seq tracks for each of these three sarcoma subtypes.

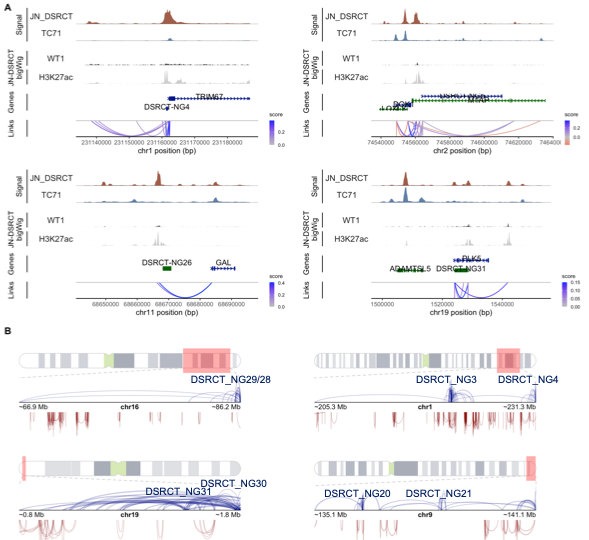

**Supplementary Figure 7. Enhancer-promoter hubs at DSRCT_NG loci promote expression of nearby genes through looping interaction. A)** Peak-to-gene associations inferred from SHARE-seq for DSRCT_NGs. Many associations originate from NG TSSs and are marked by EWS::WT1 and H3K27ac. **B)** Loop distances involving DSRCT_NGs (dark blue) and non-EWS::WT1 binding sites (dark red). Loops with DSRCT_NGs show longer interaction distances.

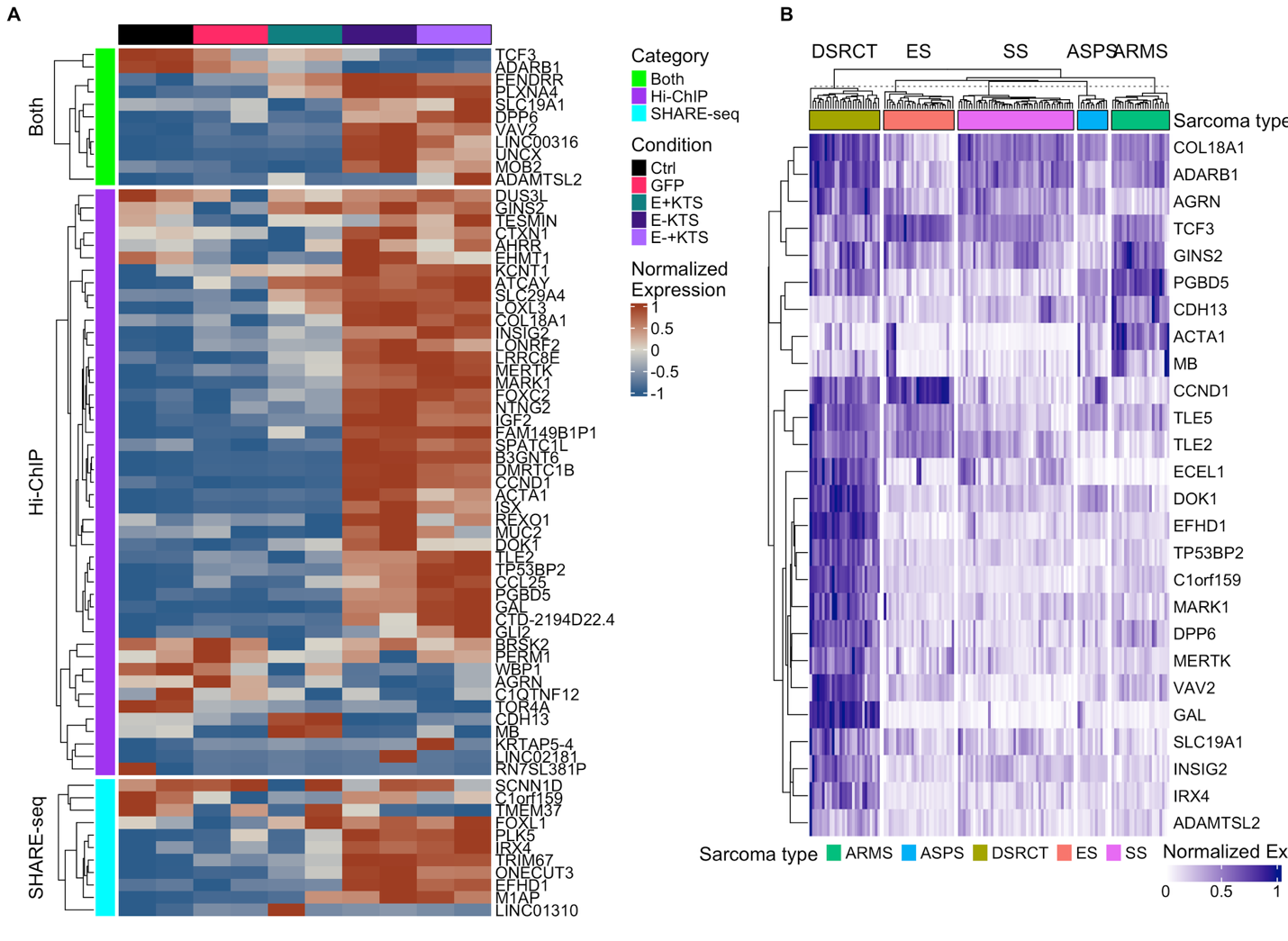
**Supplementary Figure 8. Expression of genes near DSRCT_NG loci. A)** Gene expression changes linked to DSRCT_NGs following EWS::WT1 isoform transduction in LP9 cells. E–KTS drives most gene expression changes. **B**) Microarray data expression of genes near DSRCT_NGs in other sarcomas driven by a fusion protein. Majority of gene expressed are specific to DSRCT.
Supplemental Table Captions (data provided in separate files)

**Supplementary Table 1.**  Clinical and molecular annotation of DSRCT specimens in this study. Supplementary Table 1: Clinical and molecular characteristics of DSRCT specimens included in this study. Annotations include clinical features and molecular profiling. Abbreviations: Y = Yes; N = No; – = Not available or unknown.

**Supplementary Table 2.** Clinical data and metadata associated with patient-derived xenograft (PDX) models were used in this study.

**Supplementary Table 3.** HOMER motif enrichment analysis of ±1 kb regions around DSRCT_NG TSSs. **A)** Table of known motifs and **B)** table of de novo motifs.

**Supplementary Table 4. A)** List of significant peak-to-gene associations within 50 kb of DSRCT_NG TSSs identified by SHARE-seq (p < 0.05). **B)** Distance between DSRCT_NG and nearby genes.

**Supplementary Table 5. A)** Loop distances involving DSRCT_NGs and EWS::WT1 binding sites, stratified by loop composition. **B)** Loop counts per promoter region for DSRCT_NGs and **C)** typical EWS::WT1 targets. **D)** Permutation test results showing overlap of DSRCT_NG promoter regions and **E)** linked genes with EWS::WT1 binding sites.
